## Supplementary Information for "CellMixS: quantifying and visualizing batch effects in single cell RNA-seq data"

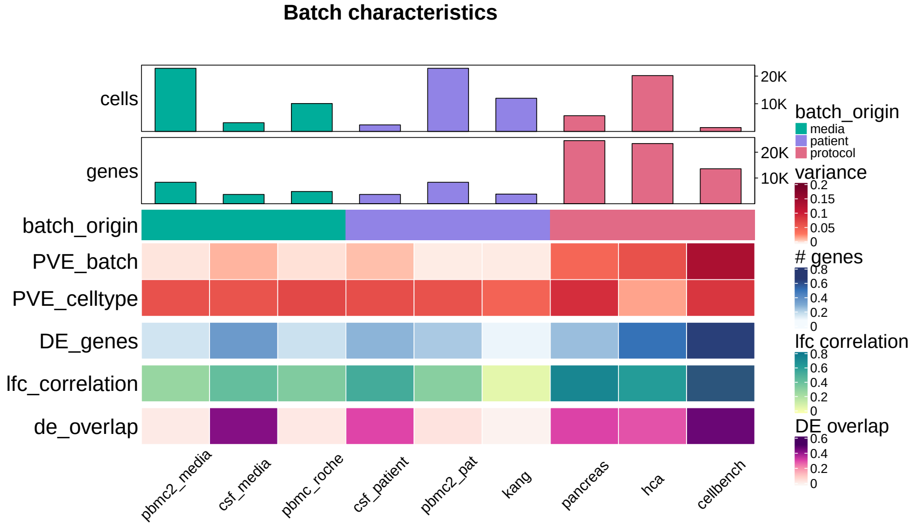


Figure S1: **Batch characterization:** Batch characteristics across 9 different batches related to differences between patients, media storage and use of sequencing protocols (batch origin). The following characteristics are shown: mean percent variance associated with the batch variable across all genes (PVE_batch), mean percent variance associated with the cell-type variable across all genes (PVE_celltype), mean relative number of differentially expressed genes between batches (DE_genes), mean Pearson correlation of log fold changes of the batch effect between cell types (lfc_correlation), mean overlap of DE genes between different cell-types (de_overlap).

| Dataset | Cells | Genes | Batches | Cluster | Access raw data | Access processed data |
|---|---|---|---|---|---|---|
| **CellBench** | 1401 | 13575 | 3 | 3 | GEO: GSE118767 | `https://bioconductor.org/packages/release/bioc/html/CellBench.html` |
| **Human Cell Atlas - Mereu** *(hca)* | 20237 | 23381 | 13 | 8 | GEO: GSE133549 | `https://www.dropbox.com/s/i8mwmyymchx8mn8/sce.all_classified.technologies.RData?dl=0` |
| **Pbmc media and patient** *(pbmc2pat, pbmc2media)* | 22824 | 8331 | 2/3 | 14 | ArrayExpress: E-MTAB-9916 | DOI: 10.6084/m9.figshare.13341200.v1 |
| **Cerebral spine fluid** *(csf_pat, csf_media)* | 3149 | 3613 | 3/2 | 5 | ArrayExpress: E-MTAB-9916 | DOI: 10.6084/m9.figshare.13341200.v1 |
| **Pbmc Roche** | 10096 | 4756 | 4 | 9 | ArrayExpress: E-MTAB-9916 | DOI: 10.6084/m9.figshare.13341200.v1 |
| **Kang** | 14619 | 3757 | 8 | 8 | GEO: GSE9658 | `https://bioconductor.org/packages/release/data/experiment/vignettes/muscData/inst/doc/muscData.html` |
| **Pancreas** | 5683 | 25381 | 3 | 13 | GEO: GSE81076, GSE85241, GSE86469, ArrayExpress: E-MTAB-5061 | `https://satijalab.org/seurat/v3.0/integration.html` `https://www.dropbox.com/s/1zxbn92y5du9pu0/pancreas_v3_files.tar.gz?dl=1` |

Table S1: **Datasets:** Short summaries on datasets included in the benchmark. Genes, cells, batches, cluster refer to the respective number per dataset. Access information to the raw and (pre-)processed data objects.

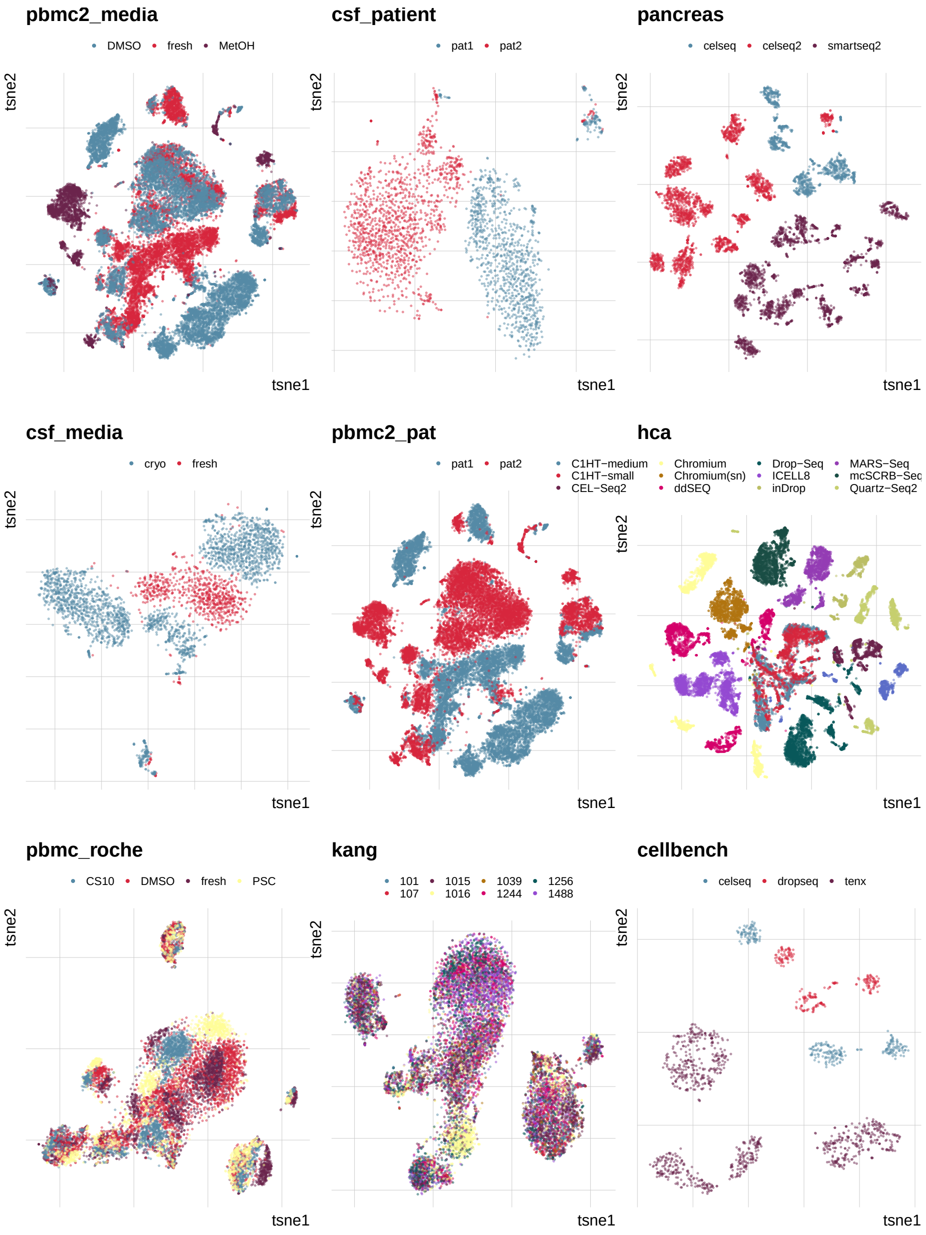


Figure S2: **Overview of batch effects and datasets included in this study:** 2D tSNE projections of batch effects characterized in this study. Different batches are indicated by colour. Batches form distinct groups in all datasets except for pbmc_roche and kang.

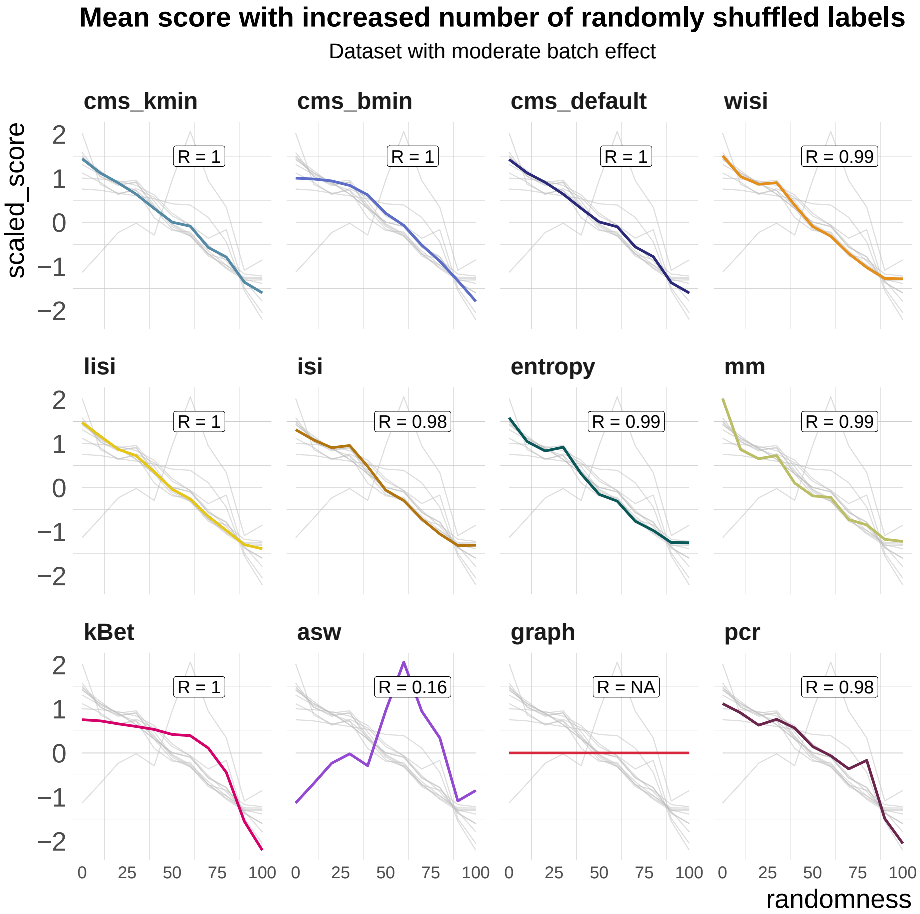


Figure S3: **Benchmark task2: Scaling with randomness:** Metric scores by increasingly randomized batch label in a dataset with a moderate batch effect (pbmc_roche). Scores were standardized by subtraction of their mean and division of their standard deviation across permutations. Directions were adjusted when necessary, such that all scores increase with batch strength. Grey lines indicate the scores of the other metrics. Corresponding absolute values of the Spearman correlation coefficients (R) are shown in the text box of each subpanel.

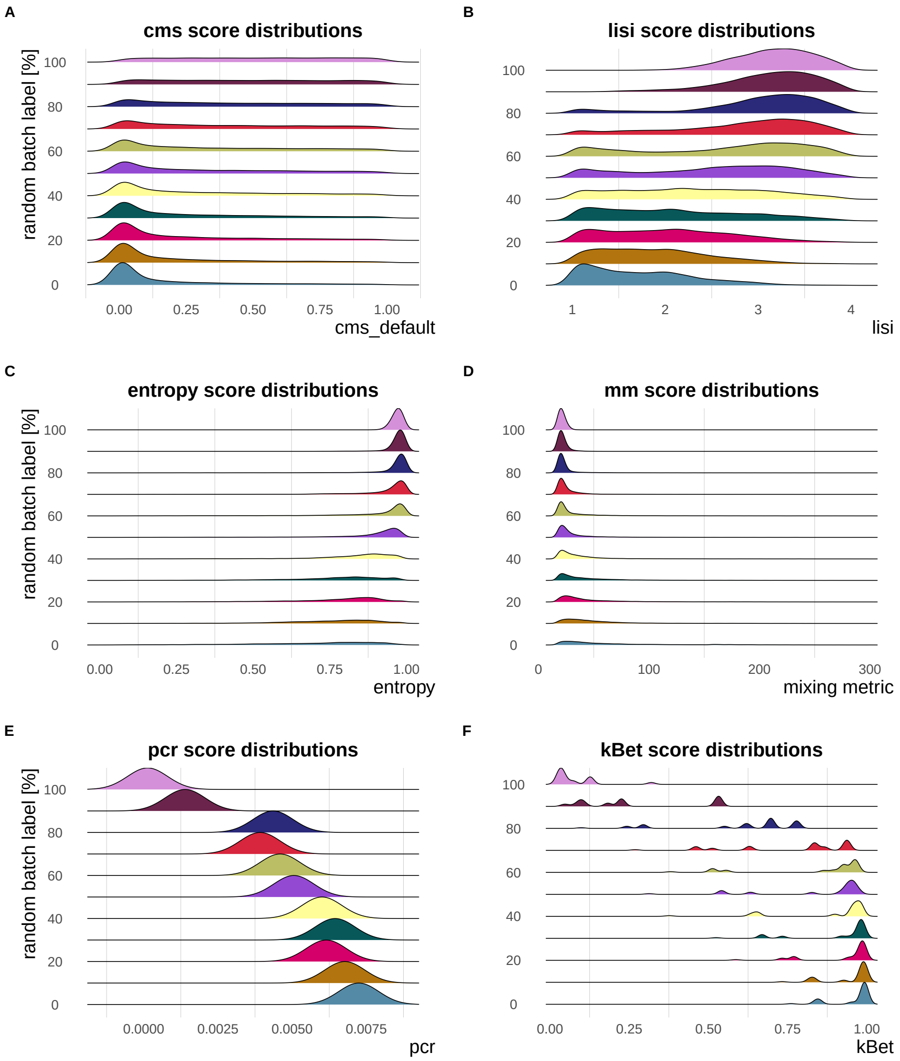


Figure S4: **Metric score distributions across data with increasingly randomized batch label:** Negative control - `cms`, `entropy`, `lisi` and `pcr` reach their nominal minimum in a dataset with completely random batch label assignments. In particular at 100% randomly permuted batch labels, `cms` shows a flat score distribution, the mean `entropy` is about 1, `lisi` approaches the effective number of batches (4 in this dataset - pbmc_roche) and `pcr` equals to 0. The mean `mm` also gradually decreases to a minimum at 100% randomly batch label permutation. While `kBet` scores decrease continuously with increasing percentage of random batch label, there are still some cell types with scores > 0 at 100% randomly assigned batch label.

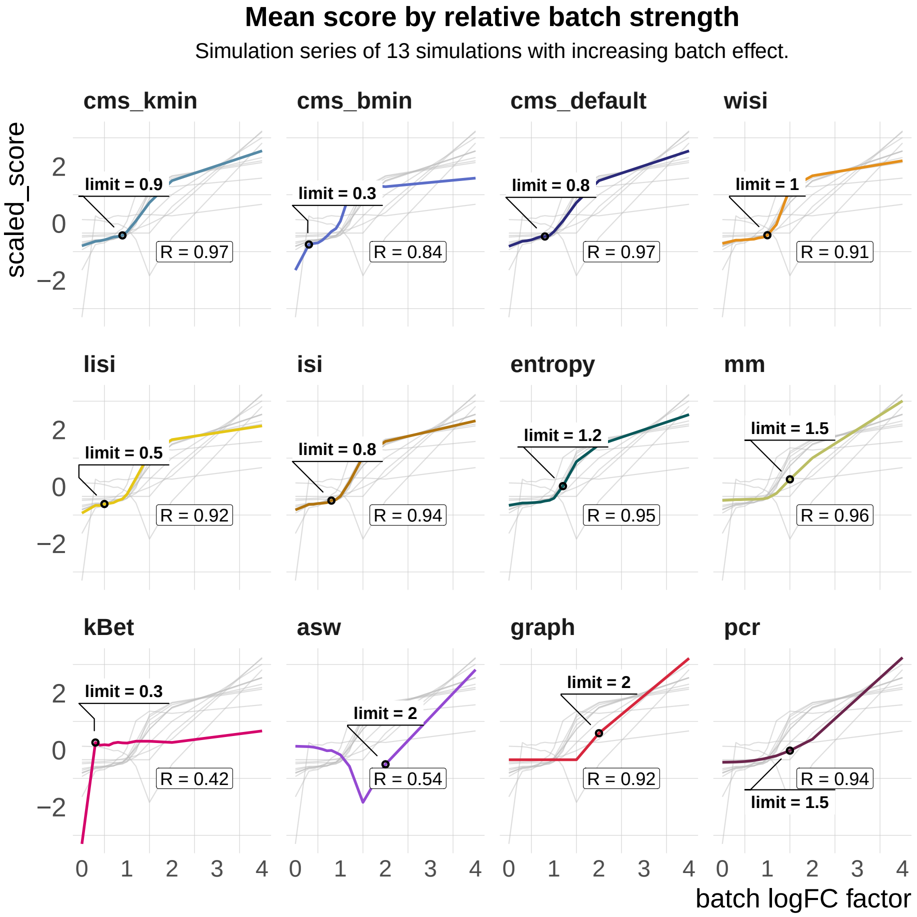


Figure S5: **Benchmark task3: Scaling and detection limits in synthetic data:** z-transformed metric scores (y-axis) across simulation series of the pbmc2_media dataset with increasing batch logFC factor (x-axis). All metrics except `asw` increase with increasing batch logFC. This is reflected in the Spearman correlation coefficient (R) between the metric's scores and the batch logFC factor. The detection limit (limit) refers to the first logFC factor with a score that differs by $\geq 10\%$ of the metric's range from the score of the batch-free simulation (batch logFC factor = 0).

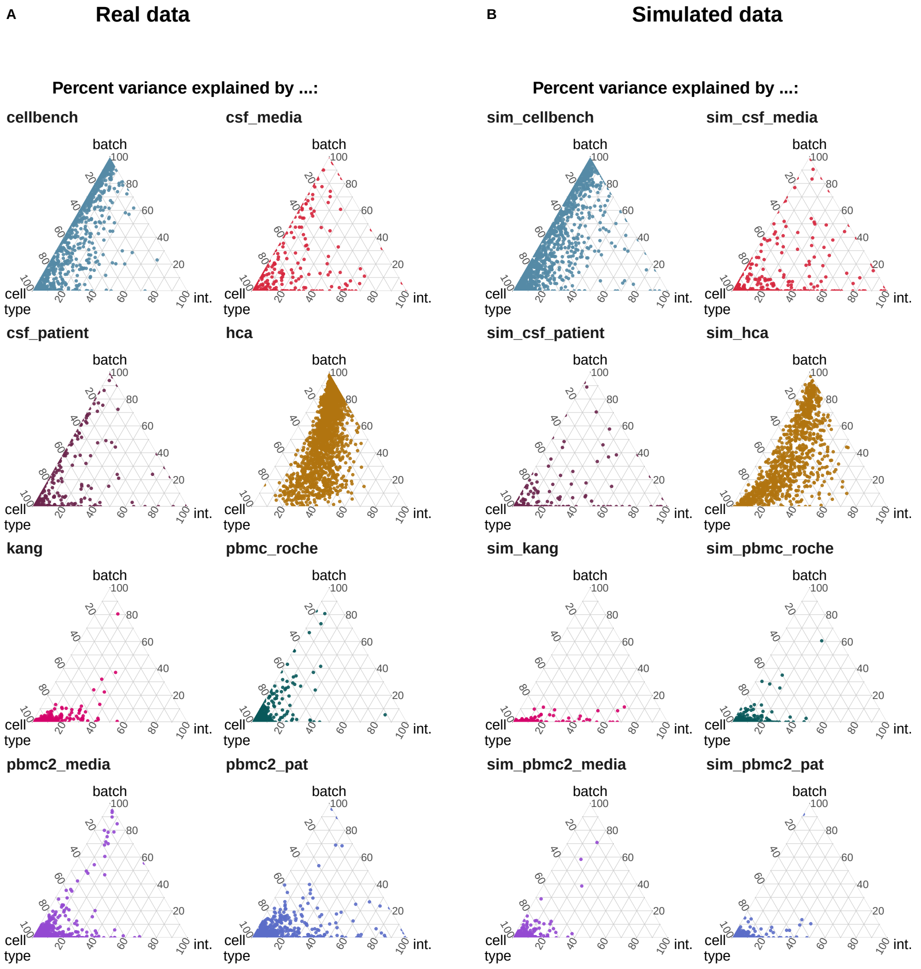


Figure S6: **Comparison of variance partitions between simulations and their corresponding real data reference**: Variance explained by batch, cell type and the interaction of cell type and batch in the real datasets (A) and the corresponding simulated datasets (B). In these simulations, the batch logFC factors are set to 1, thus simulations mimic their respective real data reference.

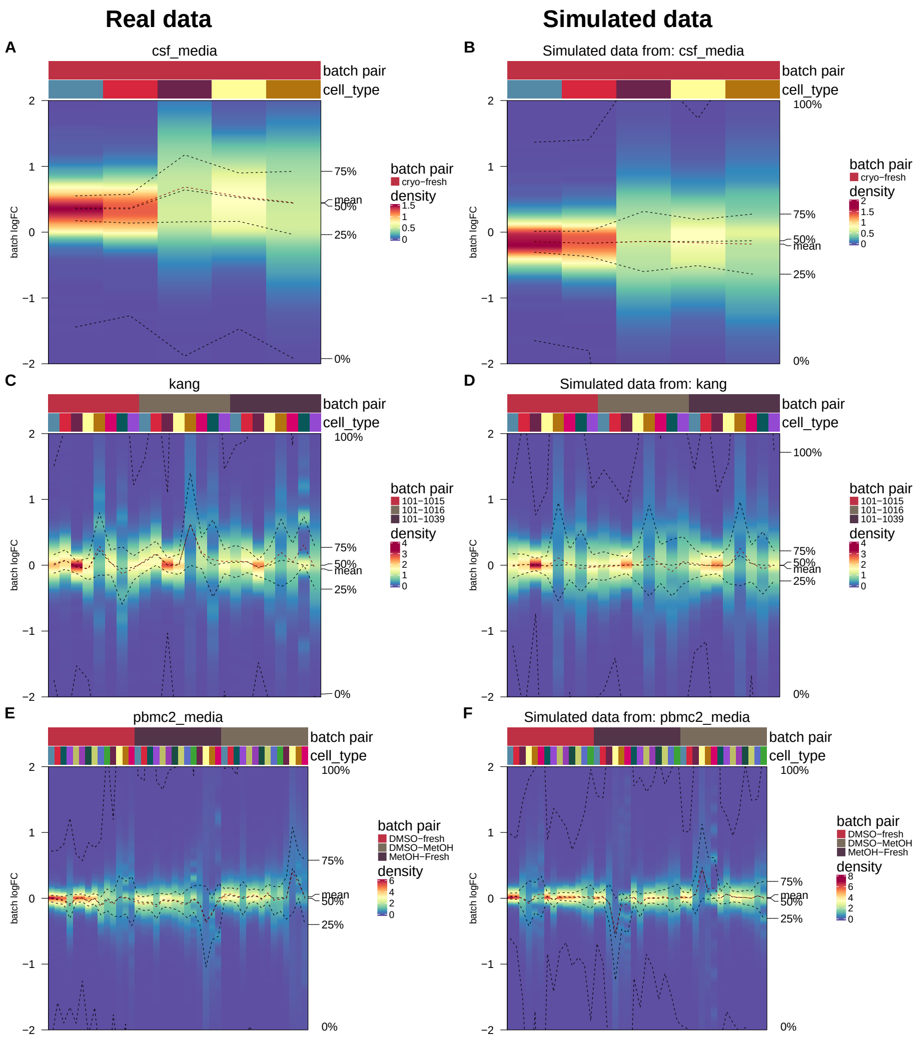


Figure S7: **Comparison of batch logFC distributions between simulations and their corresponding real data reference**: Cell type-specific logFC distributions between batches in real datasets (A,C,E) and their corresponding simulated datasets (B,D,F). Batch tuning factors are set to 1, such that simulations mimic their respective real data reference.
